# Synthetic Cooling Agent in Oral Nicotine Pouch Products Marketed as “Flavor-Ban Approved”

**DOI:** 10.1101/2023.02.23.529797

**Authors:** Sairam V. Jabba, Hanno C Erythropel, Jackson G. Woodrow, Paul T Anastas, Stephanie S. O’Malley, Suchitra Krishnan-Sarin, Julie B Zimmerman, Sven E. Jordt

## Abstract

**Background:** US sales of oral nicotine pouches (ONPs) have rapidly increased, with cool/mint-flavored ONPs the most popular. Restrictions on sales of flavored tobacco products have either been implemented or proposed by several US states and localities. Zyn, the most popular ONP brand, is marketing Zyn-”Chill” and Zyn-”Smooth” as “Flavor-Ban Approved”, probably to evade flavor bans. At present it is unclear whether these ONPs are indeed free of flavor additives that can impart pleasant sensations such as cooling.

**Methods:** Sensory cooling and irritant activities of “Flavor-Ban Approved” ONPs, Zyn-”Chill” and “Smooth”, along with “minty” varieties (Cool Mint, Peppermint, Spearmint, Menthol), were analyzed by Ca2+ microfluorimetry in HEK293 cells expressing the cold/menthol (TRPM8) or menthol/irritant receptor (TRPA1). Flavor chemical content of these ONPs was analyzed by GC/MS.

**Results:** Zyn-”Chill” ONP extracts robustly activated TRPM8, with much higher efficacy (39-53%) than the mint-flavored ONPs. In contrast, mint-flavored ONP extracts elicited stronger TRPA1 irritant receptor responses than Zyn-”Chill” extracts. Chemical analysis demonstrated the presence of WS-3, an odorless synthetic cooling agent, in Zyn-”Chill” and several other mint-flavored Zyn-ONPs

**Conclusions:** Synthetic cooling agents such as WS-3 found in ‘Flavor-Ban Approved’ Zyn-“Chill” can provide a robust cooling sensation with reduced sensory irritancy, thereby increasing product appeal and use. The label “Flavor-Ban Approved” is misleading and may implicate health benefits. Regulators need to develop effective strategies for the control of odorless sensory additives used by the industry to bypass flavor bans.

## Background

The sales of flavored oral nicotine pouch (ONP) products have rapidly increased.^1,2^ ONPs are currently marketed in a wide variety of fruit, candy, menthol/mint and concept flavors. The most popular ONP brand, Zyn (Swedish Match / PMI) is advertising two of their products, labelled “Chill” and “Smooth”, as “Flavor-Ban Approved”,^3^ likely in anticipation of regulatory restrictions on characterizing flavors in this product category. It remains unclear whether ONPs marketed as “Flavor-Ban Approved” are indeed free of flavor additives, including constituents imparting pleasant sensations such as cooling. Since the name of one of the products, Zyn-Chill, implies a cooling effect, we examined the “Flavor-Ban Approved” ONPs for sensory cooling activity and the presence of synthetic odorless cooling agents and mint flavorants, comparing with ONPs named for their mint characterizing flavors. Synthetic cooling agents such as WS-3 or WS-23 are increasingly used in electronic cigarette products, but haven’t been reported as ingredients in ONPs.^4–6^

## Methods

### ONP extract preparation

ONPs marketed as “Flavor-Ban Approved” (Zyn “Chill”, “Smooth”) and ONPs labelled for cool/mint characterizing flavors (“Cool Mint”, “Peppermint”, “Spearmint” and “Menthol”) were purchased at 3 and 6 mg nicotine / pouch strength between July and October 2022, either directly from Zyn’s official website (us.zyn.com) or from local gas stations. For calcium microfluorimetry, ONP contents were stirred overnight in calcium assay buffer (Hank’s Balanced Salt Solution with 10 mM HEPES) and dilutions of these extracts (diluted 1X-100X in assay buffer) were prepared to test for receptor activity. 1X dilution is defined as the extract of one ONP pouch in 10 mL assay buffer, and 100X the 100-fold dilution thereof. For chemical analysis, ONP contents were weighed and extracted in 2 mL of methanol for 7d in the dark with occasional stirring.

### Testing for sensory cooling and irritant activity by calcium microfluorimetry

Buffer ONP extracts (diluted 1X-100X) were tested for activity on the human cold/menthol receptor, TRPM8, and the menthol irritant receptor, TRPA1, by intracellular calcium microfluorimetry in HEK-293t cells (RRID:CVCL1926) as described.^7,8^ Responses to maximally activating L-menthol (1 mM), WS-3 (100 µM), cinnamaldehyde (1 mM), and a nicotine dilution series (1X: 6 mg/10mL, 1X-100X)) were measured for comparison.

### Chemical Analysis

Methanolic ONP extracts (in triplicate) were filtered (0.22µm) and 10 uL were injected directly into a GC/MS (Perkin-Elmer Clarus 580-SQ8S) using previously established methods.^4,8^ Extracts were analyzed for the presence of synthetic cooling agents commonly used in food products and other tobacco product categories, including WS-3 (CAS No. 39711-79-0), WS-5 (68489-14-5), WS-12 (68489-09-8), WS-23 (51115-67-4), and Frescolat MGA (63187-91-7), ML (17162-29-7), and XCool (1122460-01-8), ^4,9^ as well as for the minty flavorants menthol (89-78-1), menthone (10458-14-7), menthyl acetate (89-48-5), and carvone (2244-16-8). Identified synthetic cooling agents or minty flavorants were quantified using a GC/FID (Shimadzu 2010) using previously established methods.^4,6^

## Results

Average total pouch mass for the Zyn pouches was 405±18 mg (n=12). Zyn-Chill ONP extract robustly activated the cold/menthol receptor TRPM8, even at 20-fold dilution, indicating the presence of a cooling agent (Figure 1A). The Zyn-Chill extract exhibited a similar potency for TRPM8 activation to mint-/menthol-flavored ONP extracts (Zyn-Cool Mint, and Zyn-Peppermint), but activation was more efficacious than from extracts of Zyn-Cool Mint (39±5%), Zyn-Peppermint (53±4%) and a maximally activating menthol concentration (1mM, 84±10%) (n=3-4) (Figure 1A). The sensory irritant receptor, TRPA1 was robustly activated by mint-flavored ONP extracts (Zyn-Cool Mint and Zyn-Peppermint), yet TRPA1 activation was significantly weaker for Zyn-Chill extracts (48±13% vs. Zyn-Cool Mint and 52±8% vs. Zyn-Peppermint; n=3-4) (Figure 1B) indicating the presence of a non-irritating cooling ingredient in Zyn-Chill. Extracts from Zyn-Smooth showed weak activity on TRPM8 and TRPA1, and only at 1X dilution (i.e., no dilution of the extract).

**Figure 1:**
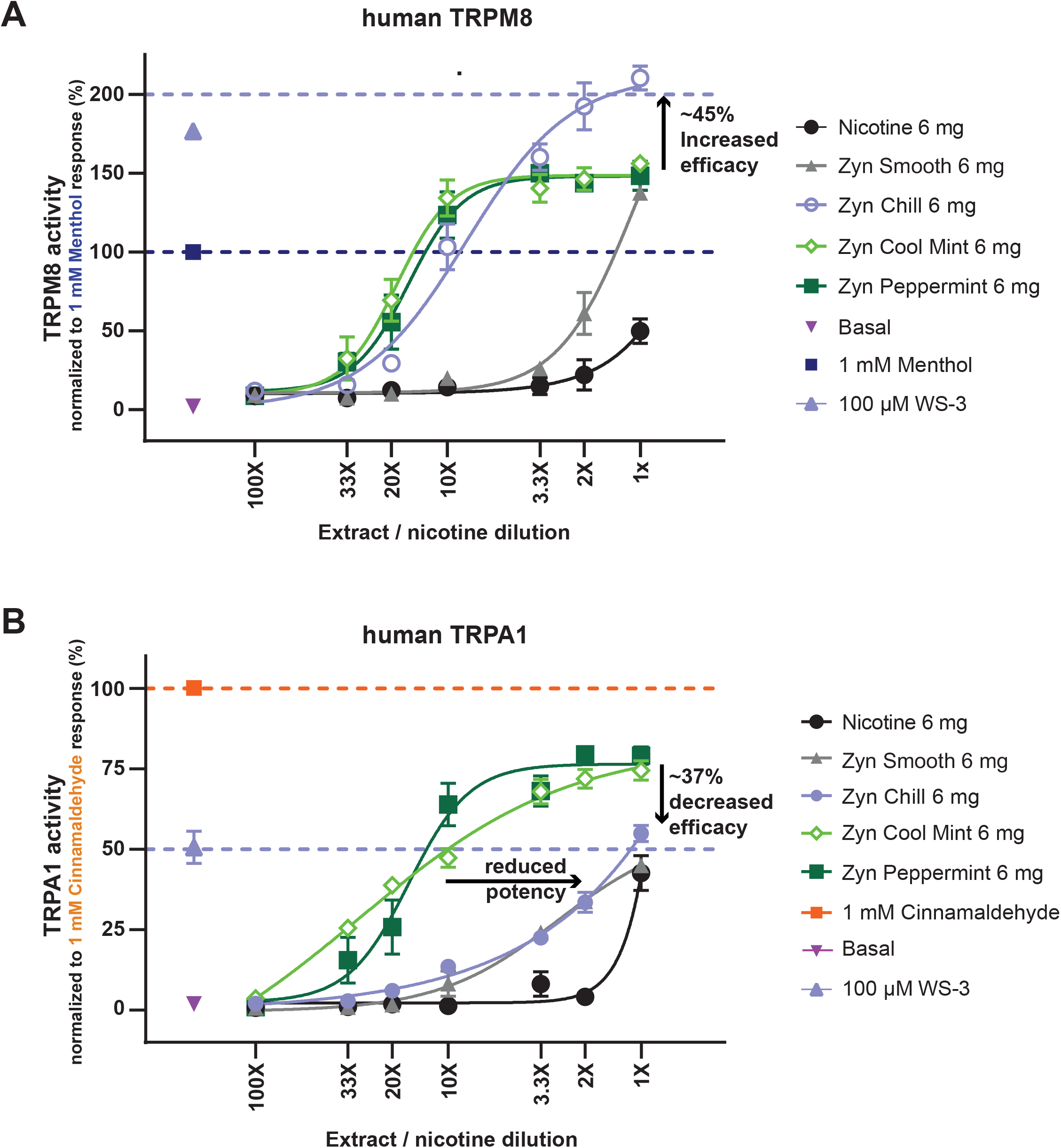
Sensory cooling and irritant activity of Zyn “Flavor-Ban Approved” and mint-flavored ONPs, measured by Ca^2+^ microfluorimetry in HEK293 cells. **A**. Dose-response analysis of human TRPM8 cold/menthol receptor-mediated Ca^2+^-influx into HEK293 cells following superfusion with a dilution series of Zyn ONP extracts (“Chill”, “Smooth”, “Cool Mint”, “Peppermint”) and nicotine (1X: 6mg/10ml). The increase in fluorescence units (F_max_-F_0_) of the fluorescent Ca^2+^ indicator (Calcium 6), was normalized to the Ca^2+^-response elicited by a saturating concentration of agonist L-menthol (1 mM). Response to a saturating concentration of WS-3 (100 µM) is shown for comparison. **B**. Dose-response analysis of human TRPA1 irritant receptor-mediated Ca^2+^-influx into HEK293 cells following superfusion with the same dilution series of ONP extracts and nicotine as in A. Ca^2+^-response to a saturating concentration of TRPA1 agonist cinnamaldehyde (1 mM) was used for normalization. Experiments were performed 3-4 times in triplicates with independent ONP extractions. Error bars for each data point show standard error of the mean.

Chemical analysis of the ONPs demonstrated the presence of WS-3, an odorless menthol-derived synthetic cooling agent in Zyn-Chill (234±7µg/pouch) as the only major flavorant and the only observable peak besides nicotine. WS-3 was also present in Zyn-Peppermint (201±11 µg/pouch) and in Zyn-Spearmint (209±15 µg/pouch). Other minty flavorants were detected in Zyn-Peppermint (Menthol: 1,925±140 µg; Menthone: 791±55 µg; and menthyl acetate: 105±9 µg) and in Zyn-Spearmint (Carvone: 1180±185 µg and Menthol: 470±44 µg). Menthol was the only flavorant present in Zyn-Cool Mint (5406±259 µg/pouch) and Zyn-Menthol (1390 µg/pouch). For Zyn-Smooth, no cooling and/or mint flavorants were detected. For WS-3 analysis, recovery was 99.5% (n=7), precision 1.8 % relative standard deviation, limit of detection 5µg/mL and limit of quantification 15µg/mL.

## Discussion

Our findings demonstrate that WS-3, an odorless synthetic cooling agent, is added to several tested cool-/mint-flavored Zyn ONPs, and to Zyn-Chill, an ONP advertised as “Flavor-Ban Approved”. Extracts from the cool-/mint-flavored ONPs also contained menthol and robustly activated the cold/menthol receptor, TRPM8. Extract from Zyn-Chill, containing only WS-3, displayed higher efficacy for TRPM8 activation, exceeding the efficacy of a maximally activating menthol concentration. While extracts of cool-/mint-flavored ONPs strongly activated the sensory irritant receptor, TRPA1, Zyn-Chill extracts showed only weak efficacy and potency at TRPA1, suggesting reduced irritancy. Containing WS-3 exclusively, “Flavor-Ban Approved” Zyn-Chill likely provides a robust oral cooling sensation by strongly activating TRPM8, in combination with lower sensory irritation (i.e. weaker TRPA1 activation) than menthol would cause, thereby likely increasing the appeal of the product.

Rodent studies demonstrated that, similar to menthol, synthetic cooling agents have analgesic activity, suggesting that WS-3 may soothe the sensory irritation caused by nicotine in ONPs.^10^ Sensory cooling stimuli, either by cold water, a synthetic cooling agent or menthol, had strong reinforcing properties in intravenous nicotine self-administration studies in rats.^11^ These studies suggest that cooling sensations and analgesia induced by odorless synthetic cooling agents in ONPs (eg., Zyn-Chill) may improve the sensory experience of the user, thereby facilitating product use initiation and more frequent consumption under the guise of a “Flavor-Ban-Approved” product.

The moniker “Flavor-Ban Approved” was likely introduced by Swedish Match to position these ONPs in markets where all flavored tobacco products have been banned, such as in the State of California.^12^ In regions without such bans, or bans that do not apply to ONPs, the products may appeal to consumers concerned about flavors associated with increased toxicity and addictiveness of tobacco products. In this case, “Flavor-Ban Approved” might represent misleading health messaging, as consumers might perceive these products as healthier. However, our present study demonstrates that these products are not free of additives modifying sensory product perception.

While the US Food and Drug Administration (FDA) has not announced any intent to regulate ONPs and their flavors, any such regulation will have to rely on the concept of “characterizing flavor” that is primarily based on odor perception, i.e., such as a minty, fruity, or sweet smell. Regulations in California and the other US states and municipalities banning certain flavored tobacco products also rely on this approach. The ambiguity of this approach may provide a loophole for the tobacco industry to add odorless sensory chemicals such as synthetic cooling agents that impart comparable or stronger physiological cooling and analgesic effects than menthol, and that at lower amounts per product unit. The manufacturer of Zyn products, Swedish Match / PMI, seems to be confident that synthetic cooling agents don’t fall under the category of “characterizing flavor” and are, therefore, “Flavor-Ban Approved.”^3^ FDA’s proposed product standard for menthol cigarettes that, if implemented, will lead to a ban of this product category, extends the definition of “characterizing flavor” by adding “multisensory experience”, including cooling sensations, as a factor to determine whether a tobacco products has a characterizing flavor.^13^ If approved as written, this extended definition will allow regulation of synthetic cooling agents in combustible cigarettes and, in the future, in all other tobacco products. Alternative approaches to regulate synthetic cooling agents include the use of positive lists of permitted tobacco additives in Canada and currently in preparation in the Netherlands,^14,15^ or Germany’s ban of chemical scaffolds present in menthol and in most synthetic cooling agents in combustible cigarettes.^16^

## Preprint Statement

This article is a Preprint and has not been peer reviewed.

## Authors Approval

All authors have seen and approved the manuscript, and it hasn’t been accepted or published elsewhere.

---

**Supplementary table 1:** Flavor chemical composition of several Zyn ONP’s. Amounts listed for each chemical are “per pouch”. n.d. – not detected

|  | Product | Flavorant (µg±stdev) |  |  |  |  |
| --- | --- | --- | --- | --- | --- | --- |
|  |  | WS-3 | Menthol | Carvone | Menthone | Menthyl acetate |
| 1 | Zyn Chill | 234±07 |  | n.d. | n.d. | n.d. |
| 2 | Zyn Peppermint | 201±11 | 1925±140 | n.d. | 791±55 | 105±09 |
| 3 | Zyn Cool Mint | n.d. | 5406±259 | n.d. | n.d. | n.d. |
| 4 | Zyn Menthol | n.d. | 1390 | n.d. | n.d. | n.d. |
| 5 | Zyn Spearmint | 209±15 | 470±44 | 1180±185 | n.d. | n.d. |
| 6 | Zyn Smooth | n.d. | n.d. | n.d. | n.d. | n.d. |

